# Modulating supplementary motor area excitability enhances groove-related pleasure during music listening

**DOI:** 10.64898/2026.05.09.722456

**Authors:** Takahide Etani, Mitsuaki Takemi, Tomohiro Samma, Jun Nitta, Saki Homma, Kenta Ueda, Keigo Yoshida, Kenjun Hayashida, Tatsuro Fujimaki, Sotaro Kondoh, Kazutoshi Kudo, Shinya Fujii

## Abstract

Pleasurable urge to move to music is often referred to as groove. Although previous studies have shown an association between the supplementary motor area (SMA) and the groove experience, its causal role remains unclear. Here, we investigated whether the SMA is causally involved in groove experience during music listening using repetitive transcranial magnetic stimulation. Fifteen healthy participants completed three sessions on separate days: excitatory stimulation (intermittent theta burst stimulation; iTBS) over the SMA, inhibitory stimulation (continuous theta burst stimulation; cTBS) over the SMA, and sham stimulation (iTBS or cTBS) over the vertex. After each stimulation session, participants listened to five high-groove and five low-groove musical excerpts and rated urge-to-move and pleasure on a 0–100 scale. Heart rate was additionally recorded as an exploratory physiological measure during music listening. Linear mixed-effects models (LMM) showed that pleasure ratings, but not urge-to-move ratings, were higher following both iTBS and cTBS compared with sham stimulation. In exploratory LMMs, reduced log-transformed heart rate variability (HRV) significantly predicted higher pleasure ratings. These findings suggest that SMA stimulation modulates the pleasurable component of the groove experience, likely via network-level mechanisms rather than a simple linear relationship between SMA excitability and pleasure. They also raise the possibility that reduced parasympathetic activity, reflected by lower HRV, mediates the stimulation-related increase in musical pleasure. Future studies should investigate the causal roles of other brain regions as well as clarify the directionality between autonomic changes and the groove experience.

## 1. Introduction

When we listen to music, there are moments when we feel a strong urge to move our bodies in time with the rhythm. This pleasurable urge to move to music is commonly referred to as groove (Etani et al. 2024; Witek et al. 2014; Senn et al. 2023; Janata et al. 2012; Vuust et al. 2022), or more recently, PLUMM (*PLeasurable Urge to Move to Music*) (Matthews et al., 2023), a term that helps distinguish it from broader definitions of groove (Duman et al., 2023). The groove experience has been shown to be influenced by multiple factors (Senn et al., 2023), including musical features such as rhythmic complexity (Matthews et al., 2019; Witek et al., 2014), beat salience (Madison et al., 2011), tempo (Etani et al., 2018; Jerjen et al., 2024), and other audio features (Stupacher et al., 2016), as well as individual differences such as musical training and personal preferences (Senn et al., 2018).

Previous studies have investigated the neural correlates of groove (Ishida et al. 2026; Stupacher et al. 2013; Matthews et al. 2020; Zampini et al. 2025; Zalta et al. 2024; Ishida and Nittono 2025), providing evidence that the groove experience is associated with brain regions involved in both reward processing and motor functions. Using transcranial magnetic stimulation (TMS), Stupacher et al., (2013) demonstrated that listening to high-groove music enhances corticospinal excitability in musicians. In a more recent study using functional magnetic resonance imaging, Matthews et al., (2020) found that high-groove rhythms increase activation in reward-related brain regions such as the nucleus accumbens and orbitofrontal cortex, as well as motor-related regions such as the supplementary motor area (SMA), premotor cortex, and basal ganglia. Additionally, using magnetoencephalography, Zalta et al., (2024) reported that beta-band oscillatory activity in the SMA is associated with groove ratings. Together, these findings suggest that motor-related brain areas, including the primary motor cortex, SMA, basal ganglia, and premotor cortex, are involved in the groove experience. This is consistent with studies on rhythm perception more generally, which show that regions such as the basal ganglia and SMA are activated during rhythm listening even when participants are instructed to remain still (e.g., Grahn & Brett, 2007).

Especially, the SMA is considered a key brain region involved in the experience of groove. As mentioned earlier, SMA activity increases when listening to rhythms or music, even in the absence of overt movement (Grahn & Brett, 2007), as well as when listening to high-groove rhythms (Matthews et al., 2020). This supports the Action Simulation for Auditory Prediction hypothesis, which proposes that beat perception is facilitated by internally simulating movement (Patel & Iversen, 2014). Within this framework, the SMA is thought to play a central role in beat processing by encoding temporal intervals and generating forward models that support rhythmic prediction (Cannon & Patel, 2021). Beyond beat perception, the SMA is also involved in temporal processing in general (e.g., Leow and Grahn 2014). Moreover, the SMA plays a role in reward processing, particularly in the anticipation of rewards (Jauhar et al., 2021). Taken together, these findings suggest that SMA activation may influence the groove experience by contributing to beat processing, time perception, and reward processing.

While the SMA has been suggested to play a role in the groove experience, its causal involvement has not been fully established. One method to examine causal relationships in brain function is repetitive transcranial magnetic stimulation (rTMS), which can modulate cortical excitability. Specifically, intermittent theta burst stimulation (iTBS) is known to increase excitability, whereas continuous theta burst stimulation (cTBS) decreases it (Huang et al., 2005). A study by Spiech et al., (2026) investigated the causal role of the SMA in the groove experience by applying cTBS to reduce its activity. In their study, participants listened to musical excerpts with low, medium, and high degrees of syncopation and rated their urge-to-move and pleasure after receiving either cTBS over the SMA or sham stimulation over the primary visual cortex (V1). Contrary to their hypothesis, urge-to-move ratings were significantly higher following the inhibitory cTBS over the SMA compared to sham stimulation, but only for rhythms with a high degree of syncopation. In contrast, pleasure ratings did not differ between stimulation conditions for any of the rhythm types. The authors speculated that reducing SMA activity may have interfered with beat perception, causing highly syncopated rhythms to be perceived as less complex and thus enhancing the groove experience.

Although a previous study has investigated the causal relationship between the SMA and the groove experience, there are several limitations. First, Spiech et al., (2026) applied only inhibitory cTBS to the SMA, without including excitatory iTBS. As a result, it remains unclear whether excitatory and inhibitory stimulation would modulate the groove experience in opposite directions, which would provide stronger evidence for a direct causal role of the SMA in the groove experience. Second, their study employed musical excerpts of 7 to 12 s long, which seem to be relatively short to assess how much groove the participants experience.

Therefore, in this study, we aimed to investigate whether modulating SMA activity influences the groove experience by comparing an excitatory stimulation condition (iTBS), an inhibitory stimulation condition (cTBS), and a sham condition. Because SMA activity has been shown to increase during the perception of high-groove rhythms (Matthews et al. 2020), we hypothesized that excitatory stimulation would enhance the groove experience (i.e., urge to move and pleasure), whereas inhibitory stimulation would reduce it, compared to sham stimulation. In addition, we aimed to explore the potential physiological effects of groove music and rTMS, as well as their interaction, by measuring participants’ heart rate, because autonomous nervous system, especially the sympathetic nervous system has been shown to be related to groove experience (Bowling et al., 2019; Spiech et al., 2022). This analysis was exploratory in nature, and no specific hypotheses were formulated.

## 2. Methods

### 2.1 Participants

Fifteen healthy individuals (mean age = 24.8 ± 7.3 years; nine females) participated in the study. The inclusion criteria were as follows: (1) no history of neurological disorders, (2) no cognitive impairments, (3) provided written informed consent to participate in the study, and (4) aged 65 years or younger on the first day of participation. The exclusion criteria were: (1) a history of seizures, (2) any medical conditions or comorbidities that could interfere with task completion, (3) any contraindications to TMS (see Supplementary Materials), (4) communication difficulties, or (5) behavior that was excessively offensive or pessimistic. The study was conducted in accordance with the Declaration of Helsinki, and the research protocol was approved by the Ethics Committee of the Faculty of Science and Technology, Keio University (IRB approval number: 2024-112). After an oral and written explanation, all participants gave an informed consent.

### 2.2 Stimuli

We selected five high-groove music and five low-groove music according to a previous study (Janata et al., 2012) (Table 1). We selected both high-groove and low-groove music because ratings for high-groove music might have been subject to a ceiling effect, such that only low-groove ratings could increase, and only high-groove ratings could decrease. We used the first 45 s of each musical excerpt.

**Table 1.** Musical excerpts used in the study (selected from Janata et al., 2012).

| High-groove music |  | Low-groove music |  |
| --- | --- | --- | --- |
| Song name | Artist | Song name | Artist |
| Flashlight | Parliament | Dawn star | Dean Magraw |
| It's a wrap | FH1 | Fortuna | Kaki King |
| Music | Leela James | Hymn for Jaco | Adrian Legg |
| Superstition | Stevie Wonder | Sweet thing | Alison Brown |
| Up for the down stroke | The Clinton Administration | Thugamar Fin | Barry Phillips |

### 2.3 Music task

Using an iPad (Apple, Inc., USA) and headphones (MDR-ZX110NC; Sony, Japan), each participant listened to ten musical excerpts 45 s each and rated the “urge-to-move” and “pleasure,” which are two main constituents of groove, using a slider (0–100). The questions used in the study were selected from the Japanese version of the Experience of Groove Questionnaire (Kawase et al., 2025; Senn et al., 2020). The musical excerpts were presented in a randomized order at each visit.

### 2.4 Heart rate recording

During the music task, heart rate was recorded using a Polar H10N sensor (Polar Electro, Japan). The Polar H10N was attached to the participant’s chest at the beginning of each experimental session.

### 2.5 Procedure

Each participant participated in three different sessions, in which they were delivered one of three TMS conditions; iTBS condition (over the left SMA), cTBS condition (over the left SMA), and sham condition (either iTBS or cTBS over the vertex) on different days. We chose to stimulate the left SMA rather than right SMA because listening to high-groove rhythms induced higher activation in the left SMA more than the right SMA in the previous study (Matthews et al., 2020). We selected the vertex as the sham condition instead of V1 used in the previous study (Spiech et al., 2026) because stimulation of V1, which is located farther from the SMA, might have made it easier for participants to notice that the stimulation was sham. After each condition, participants completed the music task. In the second or third session, they answered to the Japanese versions of the Goldsmiths Musical Sophistication Index (Gold-MSI) (Müllensiefen et al., 2014; Sadakata et al., 2022) and the Japanese version of the Barcelona Music Reward Questionnaire (BMRQ) (Mas-Herrero et al. 2014; Honda et al. 2026) after the music task. The entire experiment lasted approximately 2 h for the first session, and 1 h each for the second and third sessions.

### 2.6 TMS

Biphasic TMS pulses were delivered using a TMS stimulator (MagPro X100; MagVenture A/S, Farum, Denmark) equipped with an actively cooled figure-of-eight coil (Cool-B65; MagVenture A/S). The coil was mounted on a collaborative robot (UR3e; Universal Robots, Odense, Denmark), and participant-specific scalp surface models reconstructed from RGBD camera data were used for robotic neuronavigation, as described previously (Takemi et al., 2026).

We first determined the resting motor threshold (RMT) and then estimated the active motor threshold (AMT), which was used to set the stimulation intensity for iTBS and cTBS, from the RMT using a previously validated conversion formula (Ma et al., 2023). RMT was assessed at the hotspot for the right abductor pollicis brevis (APB) muscle, while motor evoked potentials (MEPs) were recorded from the right APB muscle using round surface electrodes (φ = 24 mm; Spes Medica, Genova, Italy). The hotspot was identified using the robotic TMS system as the scalp position that elicited the largest APB MEPs at the lowest stimulation intensity (see Takemi et al., 2026 for details). During RMT and hotspot identification, the coil was maintained tangentially to the scalp, with the handle oriented anteriorly at an angle of 45° relative to the sagittal midline, thereby inducing an initial anterior-posterior cortical current, while participants remained at rest.

TBS consisted of bursts of three pulses at 50 Hz repeated at 5 Hz (Huang et al., 2005). For iTBS, 2-s trains followed by 8 s of rest were repeated until 600 pulses had been delivered (20 cycles; 192 s in total). For cTBS, a continuous train was delivered for 40 s (600 pulses). Both iTBS and cTBS were delivered over the SMA, defined as a point 3 cm anterior and 0.5 cm left of Cz according to the international 10–20 system (Cona et al. 2017). The sham condition consisted of either iTBS or cTBS delivered over the vertex (Cz) in a counterbalanced manner across participants. The coil orientation was set at 90° for the SMA (inducing an initial medial-lateral cortical current) and at 45° for the vertex (inducing an initial anterior-posterior cortical current). Coil position was continuously tracked using a frameless stereotaxic motion capture system (V120 Duo; OptiTrack, Corvallis, OR, USA). Throughout the experiment, pulse delivery was automatically paused whenever the coil center deviated by > 4 mm or > 5° from the target position. Consistent contact between the coil and the scalp was maintained by controlling the contact pressure, measured by the collaborative robot’s internal force sensor, within a range of 0.4–1.0 kgf/cm².

### 2.7 Data analysis

For the heart rate data, prior to analysis, we excluded outliers based on the following criteria: we calculated the mean R–R interval for each participant in each 45-second trial (musical excerpt), and excluded any data points where the R–R interval was greater than 1.5 times or less than two-thirds (1/1.5) of the trial’s mean R–R interval. As an index of heart rate variability (HRV), we calculated the root mean square of successive differences (RMSSD) using the following formula:

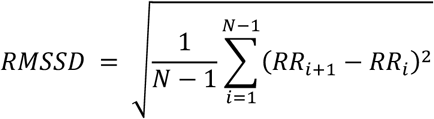

where *i* denotes the *i*th R–R interval, and *N* denotes the total number of R–R intervals in that trial. Finally, heart rate variability values were log-transformed prior to analysis to improve normality. We chose RMSSD as the measure of heart rate variability because it reflects parasympathetic nervous system activity and can be calculated from relatively short-duration recordings (Munoz et al. 2015).

### 2.8 Statistical analysis

Statistical analyses were conducted using linear mixed-effects models (LMMs) implemented in R using the *lme4* package (Bates et al. 2015). Separate LMMs were fitted for pleasure ratings, urge-to-move ratings, mean heart rate (HR), and HRV. Music condition (high-groove music vs. low-groove music), stimulation condition (iTBS vs. cTBS vs. sham), and their interaction were included as fixed effects, and participant was modeled as a random intercept to account for repeated measures. General Musical Sophistication scores of the Gold-MSI and the BMRQ total scores were included as covariates in all models. We included these factors as covariates because individuals differ in how, and to which types of music, they experience groove (Etani et al. 2025; Senn et al. 2018; Senn et al. 2023; Pando-Naude et al. 2024; Stupacher et al. 2025), and both musical ability and reward sensitivity to music have been shown to affect the groove experience (Duncan and Orgs 2024; Benson et al. 2024). Stimulation condition was treatment-coded with sham (vertex) stimulation as the reference level. Model residuals were inspected using Q–Q plots to confirm approximate normality (see Figure S1-S4). Fixed effects were evaluated using Wald tests.

Post hoc pairwise comparisons were performed using general linear hypothesis testing implemented in the *multcomp* package (Hothorn et al. 2008). Tukey correction was applied to control for multiple comparisons. When significant interactions were observed, simple-effects analyses were conducted by performing Tukey-corrected pairwise comparisons within each relevant factor level. A *p* value of < 0.05 was considered statistically significant.

For each outcome variable Y*_ijk_* (urge-to-move, pleasure, mean HR, or HRV), the following linear mixed-effects model was fitted:

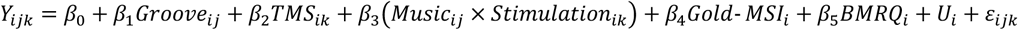

where Y*_ijk_* denotes the response of participant *i* for stimulus *j* under music condition *j* and stimulation condition *k*; β_0_–β_5_ are fixed-effect coefficients; *u_i_* represents a random intercept for participant *i* assumed to follow a normal distribution with mean zero; and ε*_ijk_* denotes the residual error term. All analyses were conducted using R (ver. 4.1.3).

## 3. Results

### 3.1 Subjective groove ratings

#### 3.1.1 Urge-to-move ratings

The results of the LMM revealed a significant main effect of music on urge-to-move ratings, with higher ratings for high-groove music compared with low-groove music (β = 12.03, SE = 2.85, *p* < 0.001) (Figure 1). In contrast, neither the main effect of stimulation condition nor the music × stimulation interaction reached statistical significance. None of the covariates showed significant effects on urge-to-move ratings. See Table S1 for detail.

**Figure 1:**
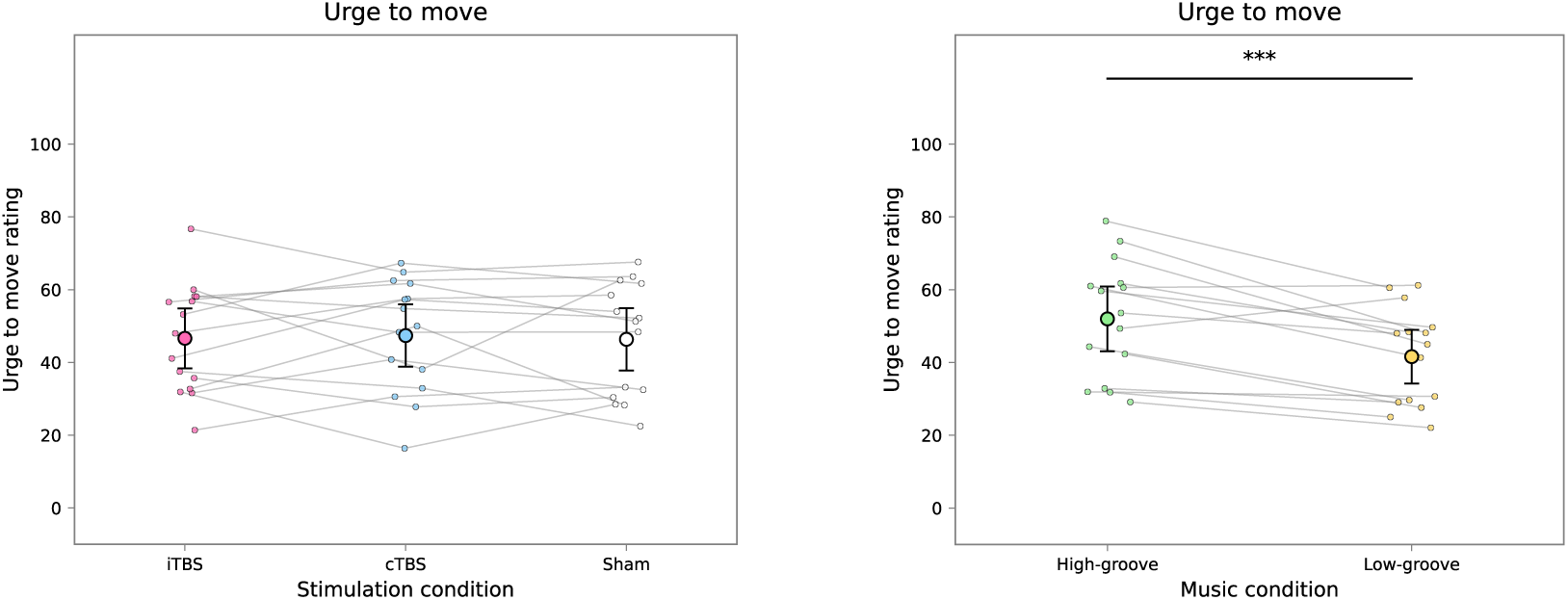
The results of the linear mixed-effects model on the urge-to-move ratings. Large points represent the observed mean values, and error bars indicate 95% confidence intervals. Small points represent each participant’s mean rating across the musical excerpts. *** *p* < 0.001

#### 3.1.2 Pleasure ratings

The LMM revealed significant main effects of stimulation and music conditions on pleasure ratings. Both iTBS and cTBS were associated with higher pleasure ratings compared with sham stimulation (iTBS vs. sham: β = 12.80, SE = 2.62, *p* < 0.001; cTBS vs. sham: β = 9.85, SE = 2.62, *p* < 0.001) (Figure 2). In addition, pleasure ratings were significantly higher for high-groove music than for low-groove music (β = 30.28, SE = 2.62, *p* < 0.001) (Figure 3). Importantly, a significant music × stimulation interaction was observed (Table S2). Post hoc simple-effects comparisons based on the fitted model indicated that, for both high- and low-groove music, pleasure ratings were significantly higher in the iTBS and cTBS conditions than in the sham condition (*p* < 0.01). No significant difference was observed between iTBS and cTBS in either music condition (Table S3). Furthermore, ratings were higher for high-groove music than for low-groove music across all stimulation conditions (Table S4).

**Figure 2:**
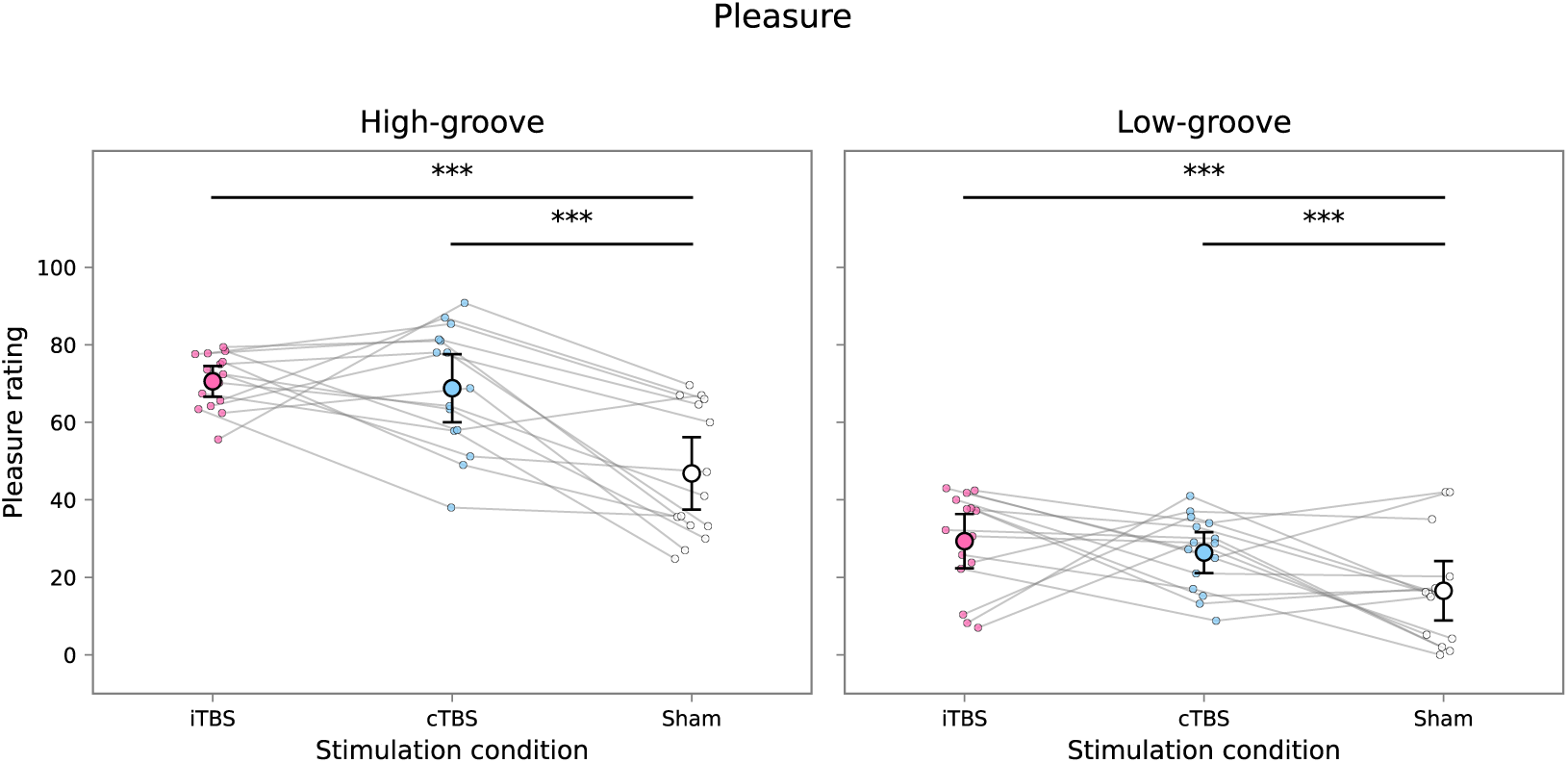
The results of the linear mixed-effects model on the pleasure ratings. Large points represent the observed mean values, and error bars indicate 95% confidence intervals. Small points represent each participant’s mean rating across the musical excerpts. *** < *p* < 0.001

**Figure 3:**
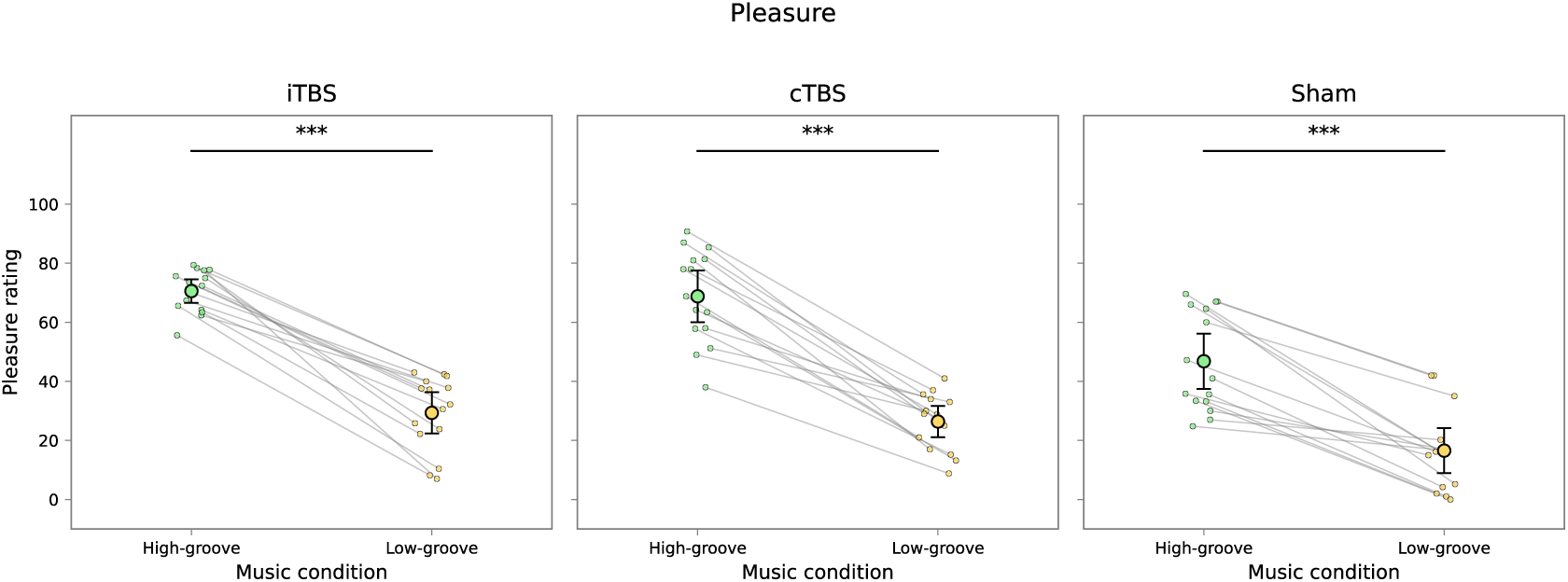
The results of the linear mixed-effects model on the pleasure ratings. Large points represent the observed mean values, and error bars indicate 95% confidence intervals. Small points represent each participant’s mean rating across the musical excerpts. *** *p* < 0.001

### 3.2 Heart rate measures

#### 3.2.1 Mean heart rate

The results of the LMM revealed a significant main effect of stimulation condition on mean HR (Figure 4). Specifically, both iTBS and cTBS were associated with significantly higher mean HR compared with sham stimulation (iTBS vs. sham: β = 3.42, SE = 0.86, *p* < 0.001; cTBS vs. sham: β = 3.43, SE = 0.86, *p* < 0.001). In contrast, the main effect of music was not significant, and no significant music × stimulation interaction was observed. The details of the results are shown in Table S5.

**Figure 4:**
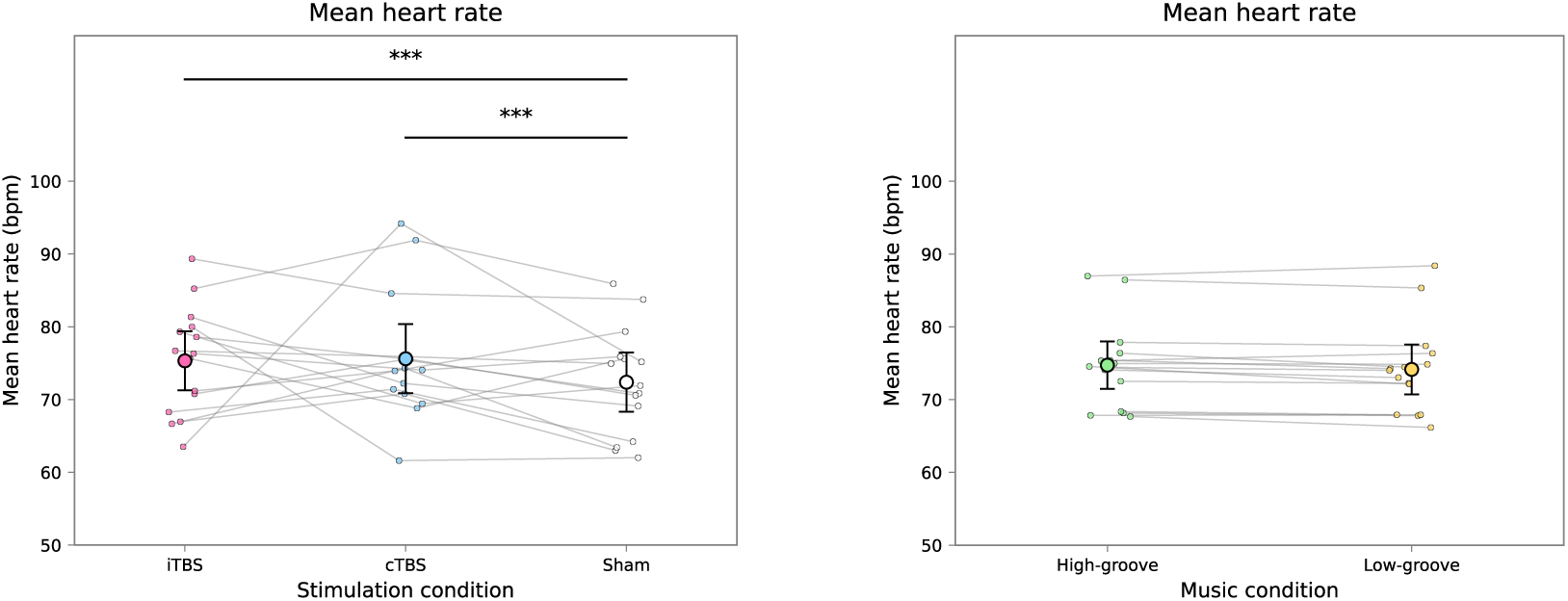
The results of the linear mixed-effects model on the mean heart rate. Large points represent the observed mean values, and error bars indicate 95% confidence intervals. Small points represent each participant’s mean rating across the musical excerpts. *** *p* < 0.001

#### 3.2.2 Heart rate variability

The LMM revealed a significant main effect of stimulation condition on log-transformed HRV (Figure 5). Specifically, both iTBS and cTBS were associated with significantly lower HRV compared with sham stimulation (iTBS vs. sham: β = −0.054, SE = 0.020, *p* = 0.0074; cTBS vs. sham: β = −0.056, SE = 0.020, *p* = 0.0052). In contrast, the main effect of music was not significant, and no significant music × stimulation interaction was observed. The details of the results are shown in Table S6.

**Figure 5:**
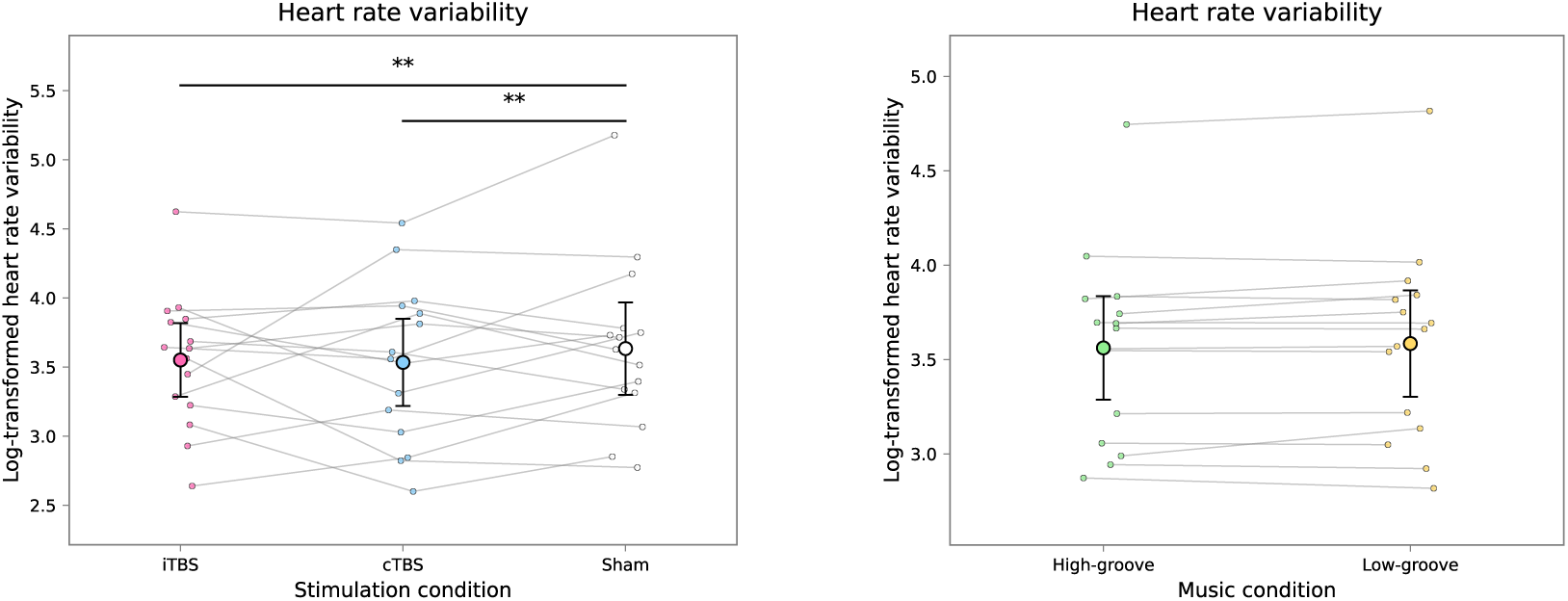
The results of the linear mixed-effects model on the mean heart rate variability. Large points represent the observed mean values, and error bars indicate 95% confidence intervals. Small points represent each participant’s mean rating across the musical excerpts. ** *p* < 0.01

### 3.3 Relationship between pleasure and hear rate measures

Because pleasure ratings, mean HR, and log-transformed HRV differed significantly between the iTBS and cTBS conditions compared with the sham condition, we explored whether pleasure ratings may be related to mean HR and HRV. To examine this, we conducted additional exploratory LMMs on pleasure ratings, adding either mean HR or log-transformed HRV as fixed effects to the original LMMs..

#### 3.3.1 Pleasure ratings and mean heart rate

Based on the LMM results, mean HR did not significantly predict pleasure ratings (β = −0.22, SE = 0.14, *p* = 0.13), whereas the main effects of music and stimulation conditions, as well as their interaction, remained significant. The details of the results are shown in Table S7.

#### 3.3.2 Pleasure ratings and heart rate variability

The LMM indicated that log-transformed HRV significantly predicted pleasure ratings (β = 7.07, SE = 13.31, *p* = 0.0020). The details of the results are shown in Table S8.

## 4. Discussion

In this study, we aimed to examine the causal relationship between the SMA activation and the groove experience applying TMS over the left SMA. In addition, we aimed to explore the potential physiological effects of musical excerpts and TBS, as well as their interaction, by measuring participants’ heart rate. The results of the LMMs revealed that while the urge-to-move ratings were not significantly affected by the TMS conditions, pleasure ratings were higher in iTBS and cTBS conditions compared to the sham condition. Our results suggest that stimulating the SMA, not dependent on excitatory or inhibitory, increases pleasure while listening to music. The results of LMMs also revealed that both urge-to-move and pleasure ratings were higher for high-groove music compared to low-groove music (Janata et al., 2012), suggesting the validity of the musical excerpts used in the study. In addition to the groove ratings, the results of LMMs revealed that the mean heart rate was higher, and the log-transformed HRV was lower for iTBS and cTBS conditions compared to the sham condition. Finally, based on these findings, we conducted additional LMMs on pleasure ratings by adding mean HR and log-transformed HRV as fixed effects. The analysis revealed that log-transformed HRV significantly predicted pleasure ratings, suggesting that HRV may mediate the relationship between music listening and experienced pleasure.

### 4.1 Urge to move

Contrary to our hypotheses, neither iTBS nor cTBS over the SMA affected urge-to-move ratings in our study. In contrast, Spiech et al., (2026) found that inhibitory stimulation of the SMA increased urge-to-move ratings for rhythms with a high degree of syncopation. Although this result was inconsistent with their original hypothesis, the authors proposed the following interpretation: because the SMA is involved in beat perception, inhibitory stimulation may have reduced the precision of beat-related predictions (i.e., lowered the precision weighting of predictive models). As a result, highly syncopated rhythms, which are normally perceived as complex, may have been perceived as less complex, thus enhancing the urge to move. In other words, beats previously perceived as off-beat may have been reinterpreted as on-beat due to the reduced precision of temporal predictions induced by SMA inhibition.

In the current study, we used musical excerpts as stimuli, selecting five high-groove and five low-groove music (Janata et al., 2012). Although we did not quantify the degree of syncopation in each excerpt, high-groove music generally tends to exhibit a medium degree of syncopation, whereas low-groove music typically contains either low or high degrees of syncopation. The low-groove excerpts used in this study lacked percussion and consisted primarily of melodic content played by instruments such as guitar, suggesting that these pieces likely had low levels of syncopation. Therefore, it is possible that the absence of highly syncopated stimuli in our experiment contributed to the lack of a positive effect of inhibitory stimulation on the SMA, as observed in (Spiech et al., 2026).

The results for excitatory stimulation may be interpreted as follows. Based on the interpretation by (Spiech et al., 2026), although excitatory stimulation may enhance the precision of rhythm perception, its effects may not be apparent when listening to stimuli with low or medium levels of syncopation. This is because such stimuli already contain a sufficient number of clearly perceived on-beats, leaving relatively few off-beat events that could be reinterpreted as on-beat through enhanced temporal precision. As a result, participants may not have experienced a stronger urge to move under excitatory stimulation in our study.

### 4.2 Pleasure

As hypothesized, pleasure ratings were higher following excitatory iTBS stimulation compared to the sham condition. However, pleasure ratings were also higher following inhibitory cTBS stimulation compared to sham condition. These findings suggest that there is no simple linear relationship between SMA activation and pleasure such as greater SMA activity leading to higher pleasure and reduced activity leading to lower pleasure. Instead, the results indicate that the SMA may influence pleasure not linearly, potentially through interactions with a broader neural network.

In Spiech et al. (2026), no significant effect of inhibitory cTBS to the SMA on pleasure ratings was observed. A key difference between their study and the present study is the length of the stimuli. While Spiech et al. (2026) used musical excerpts of 7 to 13 s long, we employed musical excerpts of 45 s long. People experience pleasure when listening to music not only from a short part but also from various information that changes throughout the entire piece including the chord progression, the change from the verse to the chorus, the break before the chorus, and the change of the key. In fact, the level of experienced pleasure during music listening changes over time (Salimpoor et al. 2009) and there are specific moments in which people experience chills when listening to music (e.g., Kondoh et al. 2026). It is therefore plausible that listening to musical excerpts long enough to experience those musical changes engages broader neural networks, including those involved in reward processing, experiencing pleasure, and motor functions. As a result, stimulating the SMA in this context may have more effectively modulated activity within these interconnected networks, including the medial orbitofrontal cortex, ventral pallidum, nucleus accumbens, and anterior cingulate cortex (Berridge & Kringelbach, 2015), ultimately enhancing the experience of pleasure.

Although the SMA is not typically considered a core region in pleasure processing, excitatory stimulation may indirectly enhance pleasure through mechanisms such as increased reward expectancy. Campos et al. (2005) found that neuronal firing in the SMA was associated with the anticipation of reward. Based on this, it is possible that excitatory stimulation of the SMA in our study may have enhanced participants’ reward expectancy during music listening, thereby amplifying the subjective experience of pleasure.

In contrast, the effect of inhibitory stimulation on the SMA may be explained by a decrease in reward sensitivity, leading to enhanced pleasure in response to smaller rewards. Vural et al. (2024) found that perturbing the pre-SMA using TMS reduced participants’ preference for larger over smaller rewards. Based on this finding, it is possible that participants in our study experienced greater pleasure from music listening, a relatively smaller or abstract reward compared to more primary rewards such as food.

### 4.3 Heart rate

Regarding heart rate, mean HR was higher and log-transformed HRV was lower in both the iTBS and cTBS conditions compared to the sham condition. As pleasure ratings were also significantly elevated in the iTBS and cTBS conditions, we conducted additional LMMs to examine whether mean HR and HRV were associated with pleasure ratings. The results revealed that HRV significantly predicted pleasure ratings, suggesting that both iTBS and cTBS may have enhanced pleasure through a reduction in HRV.

Several studies have investigated the relationship between pleasure and chills during music listening and autonomic nervous system activity, particularly heart rate responses. Listening to relaxing music has been associated with decreased heart rate, reflecting reduced sympathetic nervous system activity, and increased HRV, indicative of enhanced parasympathetic activity (Mojtabavi et al., 2020). In contrast, listening to highly arousing pleasurable music, or music that elicits chills, has been shown to increase skin conductance responses, suggesting heightened sympathetic nervous system activation (Bannister & Eerola, 2018). In the present study, participants may have experienced chill-like pleasure following iTBS and cTBS stimulation, which led to reduced HRV, indicating suppressed parasympathetic nervous system activity. This finding is consistent with a previous groove study, in which listening to high-groove music (typically rated as highly pleasurable) resulted in greater pupil dilation, which is a marker of increased sympathetic nervous system activity (Bowling et al. 2019). Taken together, our findings and those of prior study suggest that the groove experience may be accompanied by increased sympathetic activation and reduced parasympathetic activity. However, although several studies have investigated the effects of noninvasive brain stimulation, including TMS, no studies have examined its effects specifically on the SMA (Schmaußer et al. 2022). Therefore, it remains unclear whether SMA stimulation first altered HRV and subsequently influenced pleasure, warranting further investigation.

In contrast to HRV, although both iTBS and cTBS increased mean HR, heart rate itself did not significantly predict pleasure ratings. This suggests that heart rate may not directly influence the subjective experience of pleasure. Rather, it is possible that individuals experience pleasure while listening to music, and this emotional response subsequently elevates heart rate. As the current study did not investigate the causal relationship between heart rate and pleasure while listening to music, future research is needed to clarify the directionality of this association.

### 4.4 limitations

There are several limitations to the present study. First, we cannot entirely rule out the possibility that participants were aware of the differences between the two TBS conditions targeting the SMA and the sham condition targeting the vertex, given that pleasure ratings were significantly higher following both iTBS and cTBS compared to sham stimulation. However, considering the close anatomical proximity between the SMA and the vertex, this likelihood appears to be low. Additionally, the absence of significant differences in urge-to-move ratings across conditions further supports the view that participants were unlikely to have been aware of the stimulation differences.

Second, the age range of participants was limited. In the present study, most participants were in their twenties, making it difficult to generalize the findings to individuals of other age groups. Future research should address this limitation by including participants across a broader age range to examine whether the observed effects hold across the lifespan.

Third, the present study did not consider participants’ music preferences. While we included individual differences in musical sophistication and reward sensitivity by incorporating scores from the Gold-MSI and BMRQ, musical preference, which has been shown to influence the groove experience (Senn et al. 2018; Senn et al. 2021), was not assessed. Future studies should address this factor, as it may moderate the effects of stimulation or stimulus type on groove-related responses.

Finally, the causal relationship between heart rate and groove-related ratings remains unaddressed. While our results suggest that TBS over the SMA may have increased pleasure ratings via reductions in HRV, the directionality of this relationship is not clearly established. Future studies are needed to clarify whether changes in autonomic activity mediate groove experiences, or conversely, whether groove-induced pleasure modulates autonomic responses.

## 5. Conclusion

The present study demonstrated that both increasing and decreasing SMA activity enhanced pleasure aspect of groove during music listening, whereas urge-to-move ratings were not affected. These findings suggest that the SMA may contribute to the experience of musical pleasure via its involvement in broader brain networks, rather than exerting a direct linear influence on pleasure itself. Moreover, our results indicated that the increase in pleasure ratings following TBS was mediated by a decrease in HRV, raising the possibility that inhibition of parasympathetic nervous system activity may facilitate heightened pleasure during music listening. Future research is warranted to further examine the causal contributions of other brain regions to the groove experience, as well as to elucidate the relationship between autonomic nervous system activity and musical groove.

## Acknowledgement

This study was supported by JST PRESTO Grant (JPMJPR23S9), and JST COI-NEXT Grant (JPMJPF2203) awarded to S.F. and JST Moonshot R&D Grant (JPMJMS2012) awarded to M.T.

## Supplementary Materials

### Supplementary Figures

**Figure S1:**
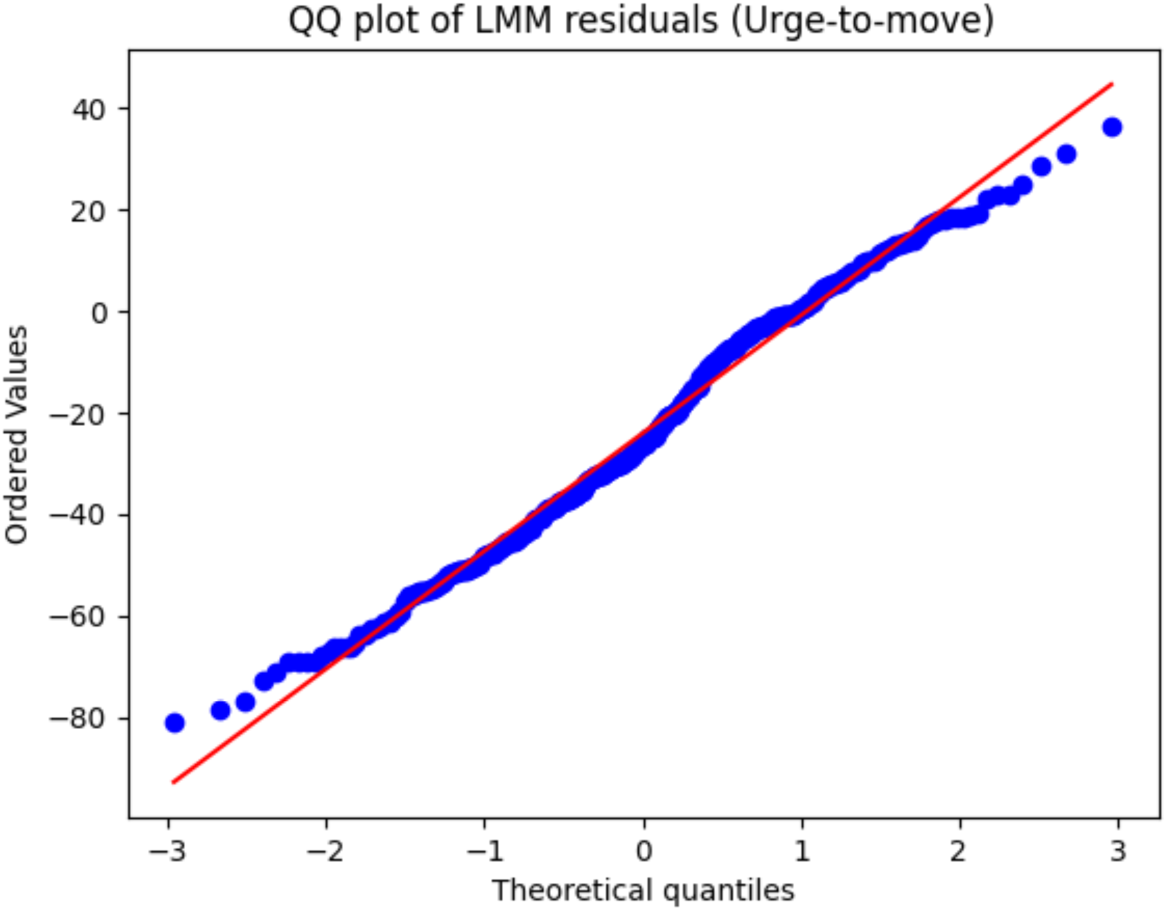
Q–Q plots of residuals from linear mixed-effects models for urge-to-move ratings. Residuals were approximately normally distributed.

**Figure S2:**
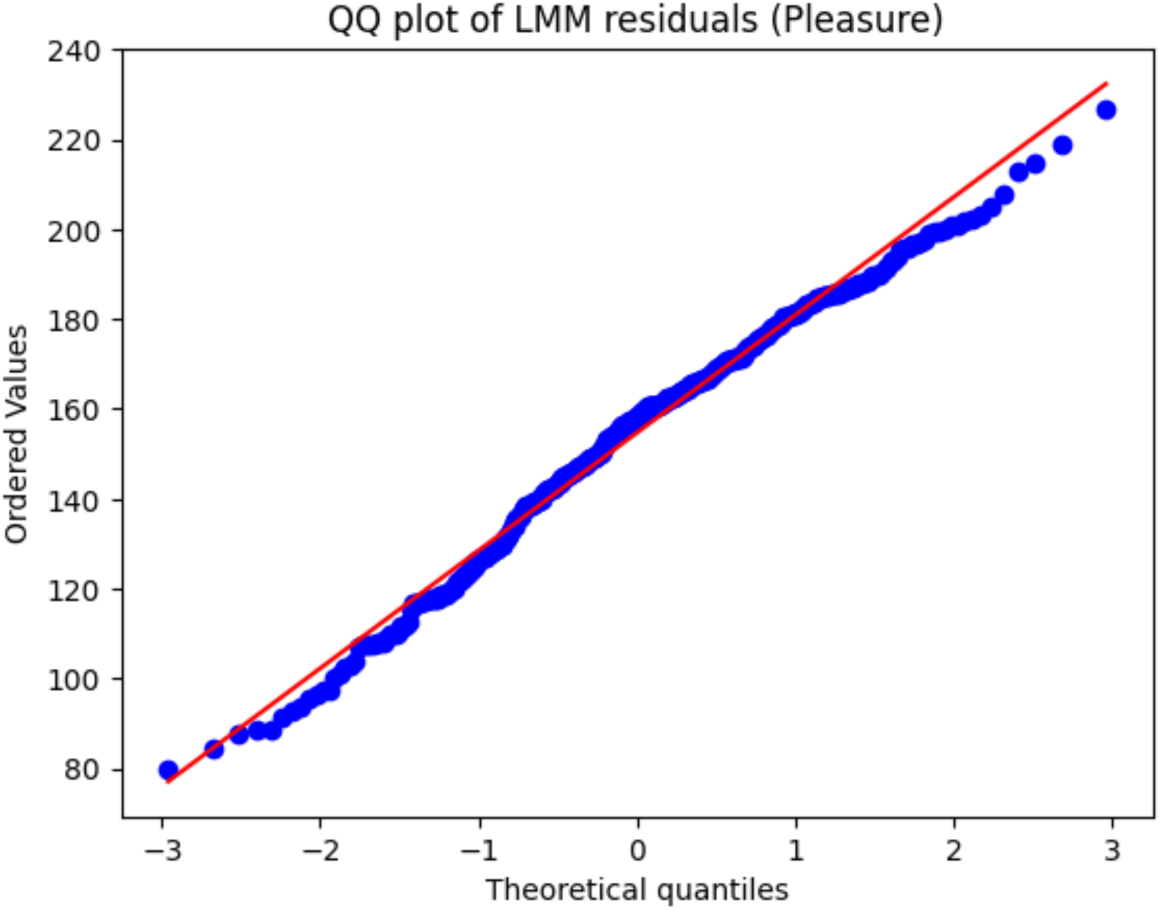
Q–Q plots of residuals from linear mixed-effects models for pleasure ratings. Residuals were approximately normally distributed.

**Figure S3:**
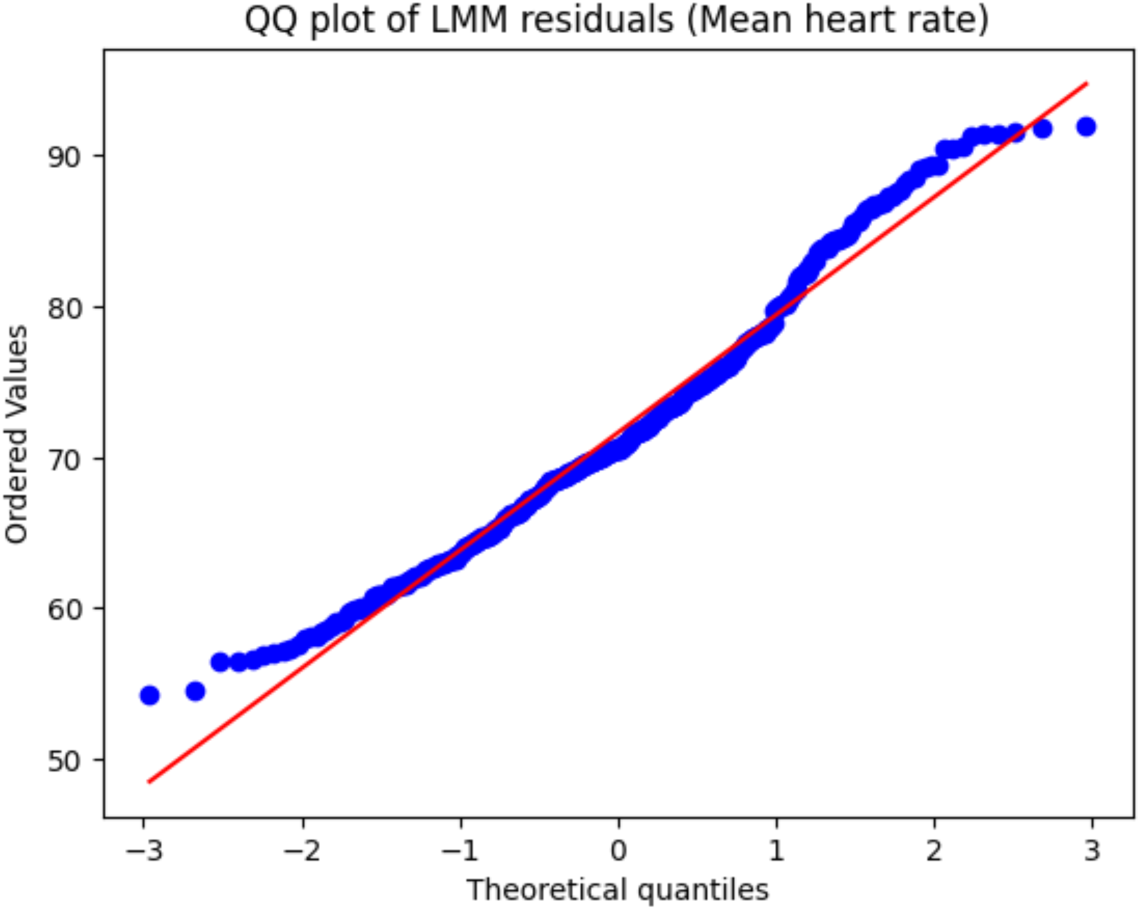
Q–Q plots of residuals from linear mixed-effects models for mean heart rate. Residuals were approximately normally distributed.

**Figure S4:**
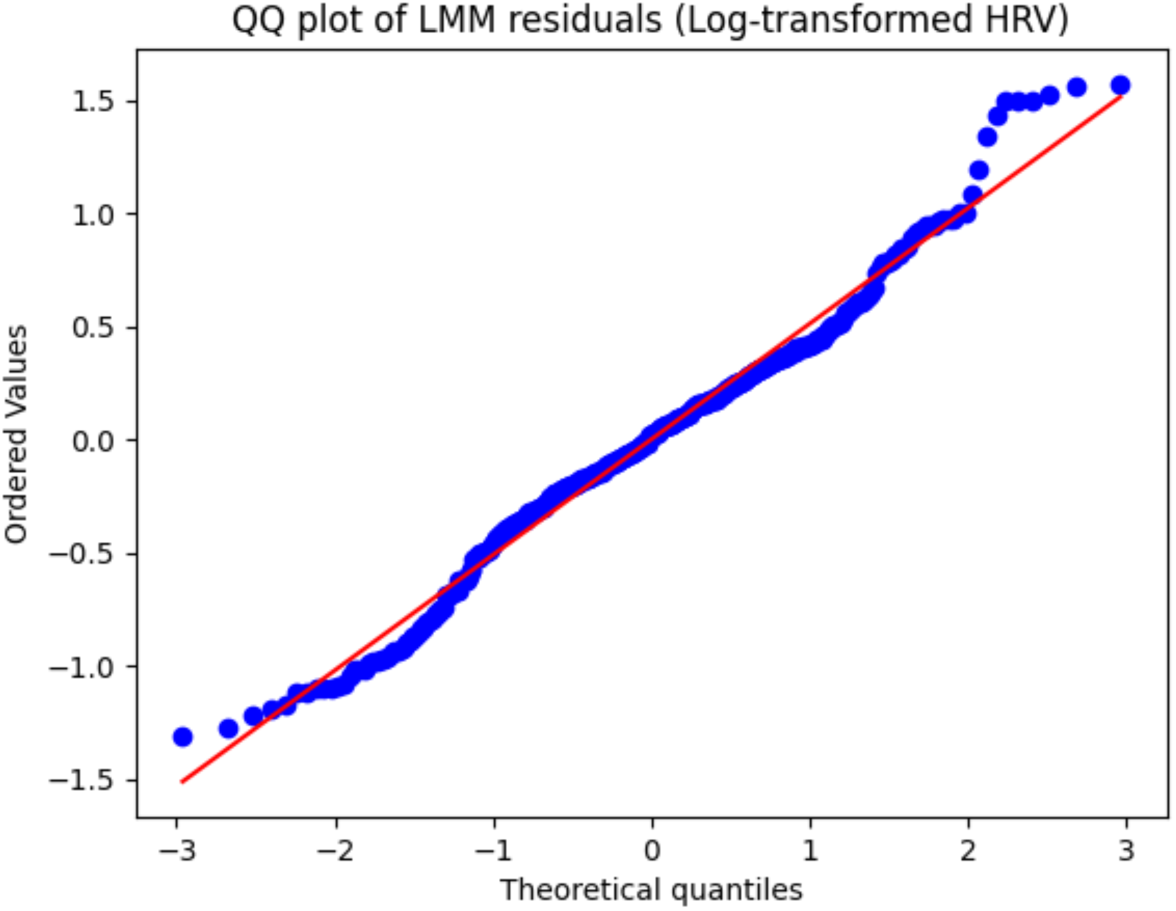
Q–Q plots of residuals from linear mixed-effects models for log-transformed heart rate variability. Residuals were approximately normally distributed.

### Supplementary Tables

**Table S1:** The results of the linear mixed-effects model on the urge-to-move ratings. *** *p* < 0.001.

| Effect | Estimate ($\beta$) | SE | t-value | p-value |
| --- | --- | --- | --- | --- |
| Intercept | 36.74 | 25.27 | 1.45 | 0.17 |
| Music | 9.51 | 2.84 | 3.35 | 0.00087*** |
| Stimulation (iTBS vs Sham) | 0.25 | 2.84 | 0.090 | 0.93 |
| Stimulation (cTBS vs Sham) | – 0.23 | 2.84 | – 0.080 | 0.94 |
| Music × iTBS | 0.053 | 4.01 | 0.013 | 0.99 |
| Music × cTBS | 2.52 | 4.01 | 0.63 | 0.53 |
| Gold-MSI General Factor | – 0.29 | 0.61 | – 0.48 | 0.64 |
| BMRQ Total Score | 0.34 | 0.47 | 0.73 | 0.48 |

**Table S2:** The results of the linear mixed-effects model on the pleasure ratings. ** *p* < 0.01 *** *p* < 0.001.

| Effect | Estimate ($\beta$) | SE | t-value | p-value |
| --- | --- | --- | --- | --- |
| Intercept | 21.88 | 12.74 | 1.72 | 0.11 |
| Music | 30.28 | 2.66 | 11.37 | < 0.001*** |
| Stimulation (iTBS vs Sham) | 12.80 | 2.66 | 4.81 | < 0.001*** |
| Stimulation (cTBS vs Sham) | 9.85 | 2.66 | 3.70 | < 0.001*** |
| Music × iTBS | 10.96 | 3.77 | 2.91 | 0.0038** |
| Music × cTBS | 12.13 | 3.77 | 3.22 | 0.0014** |
| Gold-MSI General Factor | 0.026 | 0.30 | 0.086 | 0.93 |
| BMRQ Total Score | – 0.092 | 0.24 | – 0.039 | 0.71 |

**Table S3:** The results of the post hoc analyses across groove conditions. *** *p* < 0.001.

| Music | Contrast | Estimate ($\beta$) | SE | z value | p-value |
| --- | --- | --- | --- | --- | --- |
| High groove | iTBS – Sham | 23.76 | 2.78 | 8.54 | < 0.001*** |
| High groove | cTBS – Sham | 21.99 | 2.78 | 7.90 | < 0.001*** |
| High groove | cTBS – iTBS | 1.77 | 2.78 | 0.64 | 0.80 |
| Low groove | iTBS – Sham | 12.80 | 2.32 | 5.52 | < 0.001*** |
| Low groove | cTBS – Sham | 9.85 | 2.32 | 4.25 | < 0.001*** |
| Low groove | cTBS – iTBS | 2.95 | 2.32 | 1.27 | 0.41 |

**Table S4:** The results of the post hoc analyses across stimulation conditions. *** *p* < 0.001.

| Stimulation | Contrast | Estimate ($\beta$) | SE | z value | p-value |
| --- | --- | --- | --- | --- | --- |
| iTBS | High – Low | 41.24 | 2.24 | 18.45 | < 0.001*** |
| cTBS | High – Low | 42.41 | 2.44 | 17.41 | < 0.001*** |
| Sham | High – Low | 30.28 | 2.43 | 12.46 | < 0.001*** |

**Table S5:** The results of the linear mixed-effects model on the mean heart rate. *** *p* < 0.001.

| Effect | Estimate ($\beta$) | SE | t-value | p-value |
| --- | --- | --- | --- | --- |
| Intercept | 88.27 | 9.39 | 9.41 | < 0.001*** |
| Music | 1.04 | 0.87 | 1.19 | 0.23 |
| Stimulation (iTBS vs Sham) | 3.42 | 0.87 | 3.93 | < 0.001*** |
| Stimulation (cTBS vs Sham) | 3.43 | 0.87 | 3.94 | < 0.001*** |
| Music × iTBS | – 0.96 | 1.23 | – 0.78 | 0.44 |
| Music × cTBS | – 0.41 | 1.23 | – 0.33 | 0.74 |
| Gold-MSI General Factor | 0.064 | 0.23 | 0.29 | 0.78 |
| BMRQ Total Score | – 0.27 | 0.18 | – 1.51 | 0.15 |

**Table S6:**
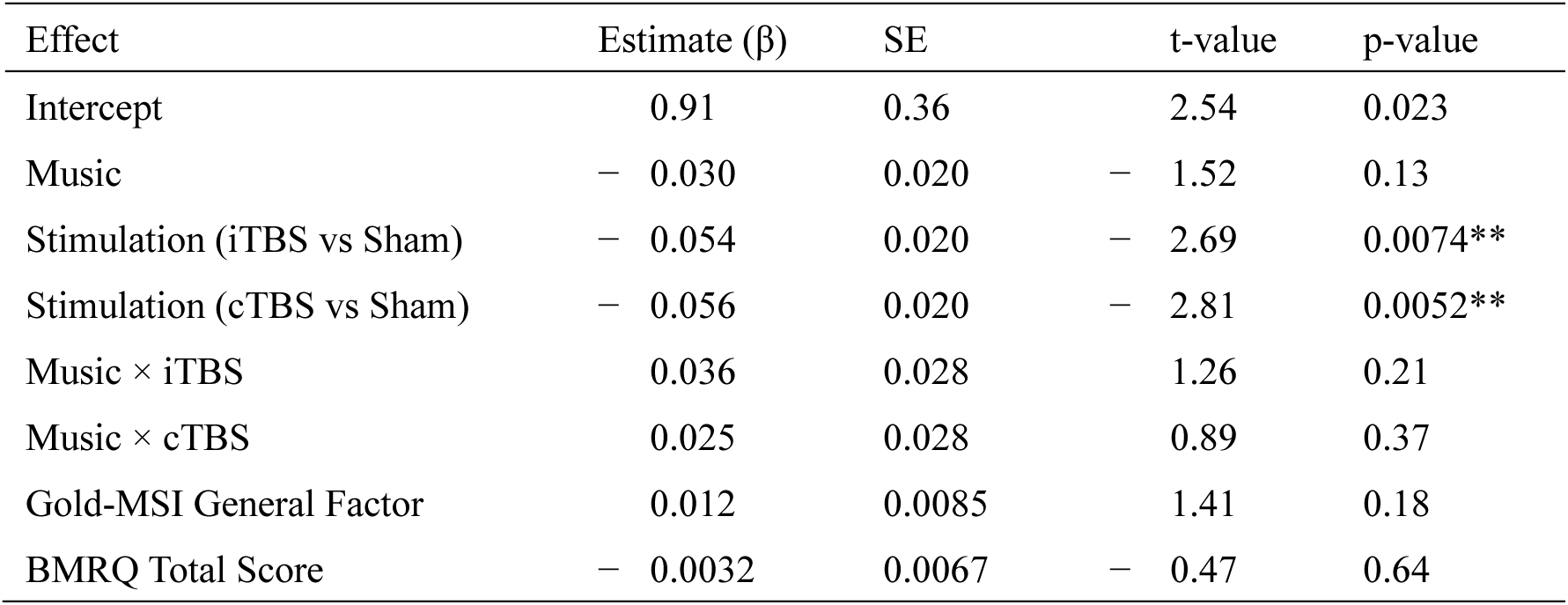
The results of the linear mixed-effects model on the log-transformed heart rate variability. ** *p* < 0.01.

| Effect | Estimate ($\beta$) | SE | t-value | p-value |
| --- | --- | --- | --- | --- |
| Intercept | 0.91 | 0.36 | 2.54 | 0.023 |
| Music | – 0.030 | 0.020 | – 1.52 | 0.13 |
| Stimulation (iTBS vs Sham) | – 0.054 | 0.020 | – 2.69 | 0.0074** |
| Stimulation (cTBS vs Sham) | – 0.056 | 0.020 | – 2.81 | 0.0052** |
| Music × iTBS | 0.036 | 0.028 | 1.26 | 0.21 |
| Music × cTBS | 0.025 | 0.028 | 0.89 | 0.37 |
| Gold-MSI General Factor | 0.012 | 0.0085 | 1.41 | 0.18 |
| BMRQ Total Score | – 0.0032 | 0.0067 | – 0.47 | 0.64 |

**Table S7:** The results of the linear mixed-effects model on the pleasure ratings adding mean heart rate as a fixed-effect. * *p* < 0.05 ** *p* < 0.01 *** *p* < 0.001.

| Effect | Estimate ($\beta$) | SE | t-value | p-value |
| --- | --- | --- | --- | --- |
| Intercept | 41.72 | 17.24 | 2.42 | 0.020* |
| Mean heart rate | – 0.22 | 0.13 | – 1.67 | 0.096 |
| Music | 30.51 | 2.66 | 11.47 | < 0.001*** |
| Stimulation (iTBS vs Sham) | 13.57 | 2.70 | 5.03 | < 0.001*** |
| Stimulation (cTBS vs Sham) | 10.62 | 2.70 | 3.94 | < 0.001*** |
| Music × iTBS | 10.74 | 3.76 | 2.86 | 0.0045** |
| Music × cTBS | 12.04 | 3.76 | 3.21 | 0.0014** |
| Gold-MSI General Factor | 0.040 | 0.30 | 0.14 | 0.89 |
| BMRQ Total Score | – 0.15 | 0.24 | – 0.64 | 0.53 |

**Table S8:** The results of the linear mixed-effects model on the pleasure ratings adding log-transformed heart rate variability as a fixed-effect. ** *p* < 0.01 *** *p* < 0.001.

| Effect | Estimate ($\beta$) | SE | t-value | p-value |
| --- | --- | --- | --- | --- |
| Intercept | 7.07 | 13.31 | 0.53 | 0.60 |
| Heart rate variability | 16.36 | 5.17 | 3.17 | 0.0020** |
| Music | 30.77 | 2.64 | 11.66 | < 0.001*** |
| Stimulation (iTBS vs Sham) | 13.68 | 2.65 | 5.16 | < 0.001*** |
| Stimulation (cTBS vs Sham) | 10.77 | 2.65 | 4.06 | < 0.001*** |
| Music × iTBS | 10.38 | 3.73 | 2.78 | 0.0056** |
| Music × cTBS | 11.72 | 3.73 | 3.14 | 0.0018** |
| Gold-MSI General Factor | – 0.17 | 0.30 | – 0.57 | 0.58 |
| BMRQ Total Score | – 0.040 | 0.23 | – 0.17 | 0.87 |

